# Electroanatomical Adaptations in the Guinea Pig Heart from Neonatal to Adulthood

**DOI:** 10.1101/2024.01.26.577234

**Authors:** Kazi T. Haq, Kate McLean, Shatha Salameh, Luther Swift, Nikki Gillum Posnack

## Abstract

**Background:** Electroanatomical adaptations during the neonatal to adult phase have not been comprehensively studied in preclinical animal models. To explore the impact of age as a biological variable on cardiac electrophysiology, we employed neonatal and adult guinea pigs, which are a recognized animal model for developmental research.

**Methods:** Healthy guinea pigs were categorized into three age groups (neonates, n=10; younger adults, n=13; and older adults, n=26). Electrocardiogram (ECG) recordings were collected *in vivo* from anesthetized animals (2–3% isoflurane). A Langendorff-perfusion system was employed for optical assessment of epicardial action potentials and calcium transients, using intact excised heart preparations. Optical data sets were analyzed and metric maps were constructed using Kairosight 3.0.

**Results:** The allometric relationship between heart weight and body weight diminishes with age, as it is strongest at the neonatal stage (R^2^ = 0.84) and completely abolished in older adults (R^2^ = 1E-06). Neonatal hearts exhibit circular activation waveforms, while adults show prototypical elliptical shapes. Neonatal conduction velocity (40.6±4.0 cm/s) is slower than adults (younger adults: 61.6±9.3 cm/s; older adults: 53.6±9.2 cm/s). Neonatal hearts have a longer action potential duration (APD) and exhibit regional heterogeneity (left apex; APD30: 68.6±5.6 ms, left basal; APD30: 62.8±3.6), which was absent in adult epicardium. With dynamic pacing, neonatal hearts exhibit a flatter APD restitution slope (APD70: 0.29±0.04) compared to older adults (0.49±0.04). Similar restitution characteristics are observed with extrasystolic pacing, with a flatter slope in neonatal hearts (APD70: 0.54±0.1) compared to adults (Younger adults: 0.85±0.4; Older adults: 0.95±0.7). Finally, neonatal hearts display unidirectional excitation-contraction coupling, while adults exhibit bidirectionality.

**Conclusion:** The transition from neonatal to adulthood in guinea pig hearts is characterized by transient changes in electroanatomic properties. Age-specific patterns can influence cardiac physiology, pathology, and therapies for cardiovascular diseases. Understanding postnatal heart development is crucial to evaluating therapeutic eligibility, safety, and efficacy.

**What is Known:** Age-specific cardiac electroanatomical characteristics have been documented in humans and some preclinical animal models. These age-specific patterns can influence cardiac physiology, pathology, and therapies for cardiovascular diseases.

**What the Study Adds:** Cardiac electroanatomical characteristics are age-specific in guinea pigs, a well-known preclinical model for developmental studies. Age-dependent adaptations in cardiac electrophysiology are readily observed in the electrocardiogram recordings and via optical mapping of epicardial action potentials and calcium transients. Our findings reveal unique activation and repolarization characteristics between neonatal and adult animals.

**Graphical Abstract:** Age-dependent adaptations in the guinea pig heart include adjustments in allometric scaling and cardiac electrophysiology.

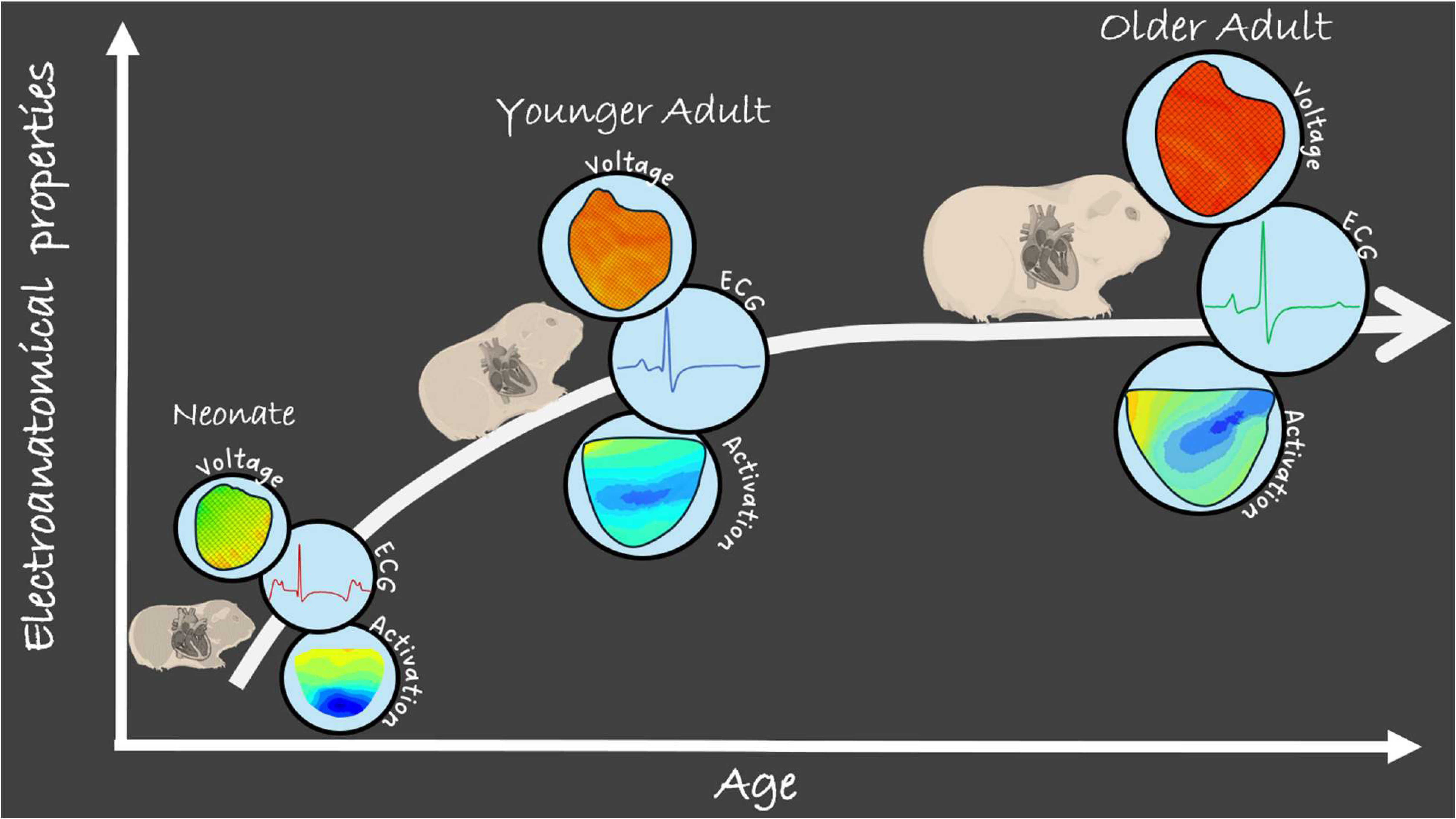

## Introduction

The scientific research community has highlighted significant challenges in precisely predicting the safety and efficacy of pediatric cardiovascular therapeutics, as preclinical models and clinical trials largely consist of adult participants^1^. As an illustrative example, milrinone, an inotropic drug commonly employed in clinical practice to enhance low cardiac output, exhibits favorable cardiovascular outcomes in adults^2^ but fails to demonstrate comparable improvements in the pediatric population^3^. This discrepancy may be attributed to distinct features in cardiovascular physiology or pathophysiology in pediatric patients, which can vary even between age groups within the pediatric population ^4^.

The cardiovascular system undergoes remarkable transformations from the neonatal period to adulthood, encompassing variations in heart rate^5^, cardiac output^4^, vascular resistance^6^, myocardial contractility^7^, and autonomic regulation^8^. For the most part, the neonatal heart functions near-maximal contractility with a heavy reliance on heart rate to modulate cardiac output, highlighting its limited cardiac reserve^9^. Conversely, the adult heart has a more dynamic response on the Frank-Starling curve, possesses greater cardiac reserve, and demonstrates heightened responsiveness to autonomic regulation^10^. At the cellular level, neonatal cardiomyocytes differ from adult cardiomyocytes in several key aspects^11^. Neonatal cardiomyocytes are smaller and maintain their proliferative capacity, increasing in total cell number until approximately six months postnatally^9^. Neonatal cardiomyocytes exhibit fewer myofibrils and mitochondria, lower intracellular calcium levels, and greater dependence on trans-sarcolemmal calcium flux^12,13^. During postnatal development, cardiomyocytes gradually acquire an adult cell morphology and functional properties, adding new sarcomeres, aligning myofibrils, and developing mature structures like T-tubules, sarcoplasmic reticulum (SR), and intercalated discs^14^.

Cardiovascular pathology also exhibits distinct variations in prevalence and prognosis across different age groups. For instance, studies in infant patient populations indicate that complete atrioventricular block and atrioventricular reentrant arrhythmia are the most common types of arrhythmias^15,16^. In contrast, among adults, atrial fibrillation, bradyarrhythmia, and conduction system diseases exhibit the highest prevalence^17^. In the benign category of arrhythmias, the prevalence of premature atrial contractions and premature ventricular contractions is nearly identical in adults^18^, while premature atrial contractions dominate in infants^19^.

Chronological age has emerged as a pivotal biological variable that influences the mechanisms underlying cardiovascular disease and modulates the safety and efficacy of various therapeutics. Yet, there is a dearth of research on the developmental trajectory of cardiac physiology and electrophysiology in animal models that are commonly employed for preclinical studies. We previously reported on the electrophysiological characteristics that are unique to pediatric rats^20^, but it is unclear whether age-specific alterations are similar across species and whether such trends have translational significance. In this study, we performed an ambispective cohort study using guinea pigs (a well-established animal model for developmental studies^21,22^) to determine whether postnatal cardiac electrophysiology exhibits distinct and transient changes – as compared to adults. To the best of our knowledge, this is the first study to investigate age-specific variations in cardiac physiology using both *in vivo* electrocardiogram (ECG) recordings and *ex vivo* optical mapping of voltage and intracellular calcium in Langendorff-perfused intact heart preparations. We evaluated characteristic ECG parameters, action potential (AP) and calcium transient (CaT) duration, conduction velocity (CV), and AP-CaT coupling characteristics under different pacing protocols (dynamic and extrasystolic) in both neonatal and adult animals.

## Methods

### Animal model

The Institutional Animal Care and Use Committee of the Children’s Research Institute approved all animal procedures, which align with the guidelines outlined in the National Institutes of Health’s *Guide for the Care and Use of Laboratory Animals*. Experiments were performed using male and female Dunkin-Harley guinea pigs procured from two premier suppliers of laboratory animals (Hilltop Lab Animals: Pennsylvania, US; Charles River Laboratories: Quebec, CA). Animals were housed in conventional acrylic cages within the research animal facility, following standard environmental conditions including a 12-hour light/dark cycle, a temperature range of 18–25°C, and humidity levels maintained between 30% and 70%. To evaluate age-specific differences, guinea pigs (n=49) were categorized into three age groups: 1) neonates (n=10, 90-125 g, 1-2 days old), 2) young adults (n=13, 300-500 g, 1-3 months old), and 3) older adults (n=26, 750-1350 g, 18-22 months old).

### Exclusion criteria

Based on our prior work^23–26^, we established a set of exclusion criteria to minimize the impact of experimental factors on outcome measurements.

1. Animals were excluded from in-vivo ECG analyses if:

- The heart rate changed sharply within 2 minutes of recording, possibly due to inadequate anesthetic response.
- ECG recording had a poor signal-to-noise ratio.
2. Animals were excluded from heart weight analysis if the measured value was identified as an outlier using statistical analysis. An overly heavy heart weight measurement can occur if fluid is not adequately extracted from the tissue (after Langendorff-perfusion).
3. Animals were excluded from optical image analysis if:

- The signal-to-noise ratio was lower than the threshold value (∼70).
- Filtering (spatial and temporal) in post-processing significantly distorted the optical signal.
- The optical signal suffered from motion artifact.
4. Sections of an optical image were excluded from analyses if the standard deviation of action potential duration (APD) or calcium transient duration (CaD) exceeded 10 ms within a selected region of interest. The latter can occur around the edges of the heart tissue due to inadequate light illumination, or within a given tissue region due to inferior dye loading.
5. In paced (dynamic or extrasystolic) data analysis, a beat at a specific pacing frequency was excluded if there was no capture or if the number of replicates fell below three.

### *In vivo* ECG

ECG recordings were performed as previously described^24,25^. Briefly, animals were anesthetized with 2–3% isoflurane for 10 minutes. Needle electrodes were placed subcutaneously to capture ECG signals in lead I configuration. Signals were acquired using a PowerLab acquisition system and LabChart 8 software (ADInstruments: Colorado US). A high-pass filter (10 Hz, analog), low-pass filter (100 Hz, analog), and gain amplifier (1k) were implemented through a differential amplifier coupled with a 60 Hz noise eliminator (Hum Bug, Digitimer: Florida, US). The timing and depth of isoflurane anesthesia were carefully controlled to minimize variability. ECG data were analyzed using LabChart 8 software. A representative beat was constructed by averaging 10 consecutive beats, and key ECG parameters (RR interval, heart rate, P duration, PR interval, QRS duration, QT and QTc (Bazette corrected QT interval^27^), and T_peak_ -T_end_) were calculated.

### Isolated heart preparation

Intact heart isolation was performed as described previously^25^. Briefly, guinea pigs were anesthetized with 2–3% isoflurane until achieving a surgical depth, as determined by an ear pinch test. Following a thoracotomy, the heart was excised and animals were euthanized by exsanguination. The whole, intact heart was cannulated at the aorta and transferred to a temperature controlled (37°C), constant pressure (68-70 mmHg) Langendorff-perfusion system. A modified Krebs-Henseleit buffer was used for the perfusate^28^, containing (in mM) 118.0 NaCl, 3.3 KCl, 2.0 CaCl_2_, 1.2 MgSO_4_, 1.2 KH_2_PO_4_, 24.0 NaHCO_3_, 10.0 glucose, 2.0 sodium pyruvate, and 10.0 HEPES buffer. The perfusate was continuously bubbled with carbogen (95% O_2_, 5% CO_2_). Intact heart preparations were allowed to equilibrate on the perfusion system for 10-20 minutes.

### Fluorescence imaging

After the heart equilibrated to the perfusion system, 12 μM (-/-) blebbistatin was introduced to the perfusate to lower metabolic demand and minimize contractile function ^29,30^, which can introduce motion artifacts in optically-acquired signals^31^. Hearts were loaded with a calcium indicator dye (50 μg Rhod-2, AM; AAT Bioquest: California US), which was recirculated for 10 minutes. Subsequently, a potentiometric dye was loaded (62 μg RH237 AAT Bioquest). The epicardial surface was illuminated using high-powered LED light sources (Solis-525C, ThorLabs: Virginia US), equipped with excitation filters (535 ± 25 nm). Fluorescence signals from Rhod-2 were acquired using a bandpass filter (585 ± 20 nm), while those from RH237 were filtered with a long-pass filter (>710 nm). Image stacks were collected from the anterior epicardium using an Optical Mapping System (MappingLabs LtD, Oxford UK) equipped with two Prime BSI cameras (Teledyne Photometrics: Arizona US) at 400-600 frames per second.

### Optical mapping and analysis

Fluorescence signals were analyzed and optical maps were generated using *Kairosight-3.0* ^26^. Activation maps (representing the time of maximum slope of the AP upstroke (dF/dt_max_)) were generated and CV was calculated to characterize epicardial AP activation in the ventricles. During apical pacing, CV was determined from the earliest to latest activation site on an epicardial activation map. Optical maps were also generated for APD and CaD to characterize the timing of AP repolarization or CaT decay. APD was measured as the difference between activation time and specific repolarization phases (30%, 50%, and 70%). Similarly, CaD was calculated as the difference between the maximum slope of the CaT upstroke (dF/dt_max_) to specific phases of CaT decay (30%, 50%, and 70%). The latency between AP repolarization and CaT decay was also determined at each pixel, as previously described^26^. To reveal restitution characteristics, dynamic (S1-S1) and extrasystolic (S1-S2) pacing was applied to the ventricular apex using different pacing cycle lengths (PCL). Finally, the derived optical maps were subdivided into four regions of interest to investigate regional heterogeneity in electrophysiology parameters.

### Immunohistochemistry

Left ventricular tissue was fixed with 10% neutral buffered formalin, paraffin embedded, and sliced to 4-5 μm thickness. Tissue slices were deparaffinized through sequential washes with Citrisolv, 100%, 95%, and 70% ethanol, followed by distilled water. Heat-induced antigen retrieval was performed in a pressure cooker with rodent decloaker buffer (RD913, Biocare Medical). Tissue slices were blocked with 3% bovine serum albumin, incubated with anti-connexin 43 rabbit primary antibody (1:200; C6219 Sigma Aldrich) overnight at 4°C, and then incubated with a goat anti-rabbit secondary antibody (1:500; Alexa Fluor 647) for 1 hour. Samples were mounted using Fluoromount-G with DAPI (00-4959-52, Invitrogen), and images were collected using a Leica TCS SP8 microscope.

### Statistical analysis

This study involved a comparison of neonates, younger adults, and older adult animals, with results presented as means ± SEM. To assess differences between two group means, an unpaired, two-tailed Student’s t-test was employed. Three or more group comparisons were analyzed using ordinary ANOVA (equal variance) or Welch’s ANOVA (unequal variance) with multiple comparisons testing. Statistical significance was determined by a p-value less than 0.05, which is denoted in each figure by an asterisk.

## Results

### Allometric scaling of heart weight in guinea pigs

Body weight is widely recognized as a reasonable predictor of age, and the proportional relationship between heart weight and body weight is well-established^32^. Anatomical and electrical allometric scaling of the heart has been observed in numerous species^33^. Accordingly, we tested whether allometric scaling of heart weight persisted in guinea pigs across different age groups. We observed a strong (R^2^ = 0.84, n=10), modest (R^2^ = 0.54, n=13), and poor correlation (R^2^ =1E-06, n=26) between heart weight and body weight in neonates, younger adults, and older adults, respectively (**Figure 1A**). To further examine this correlation, we applied a parabolic fit to each dataset, which revealed variable slopes between age groups: highest in neonates (0.013), slightly flatter in younger adults (0.011), and approaching a flat trend in older adults (**Figure 1B**). Furthermore, this trend is consistently reflected in the heart weight-to-body weight ratio, which decreases with age (neonate: 7.61±0.86 g/Kg; younger adult: 6.80±1.44 g/Kg; older adult: 6.18±1.85 g/Kg, **Figure 1C**).

**Figure 1.**
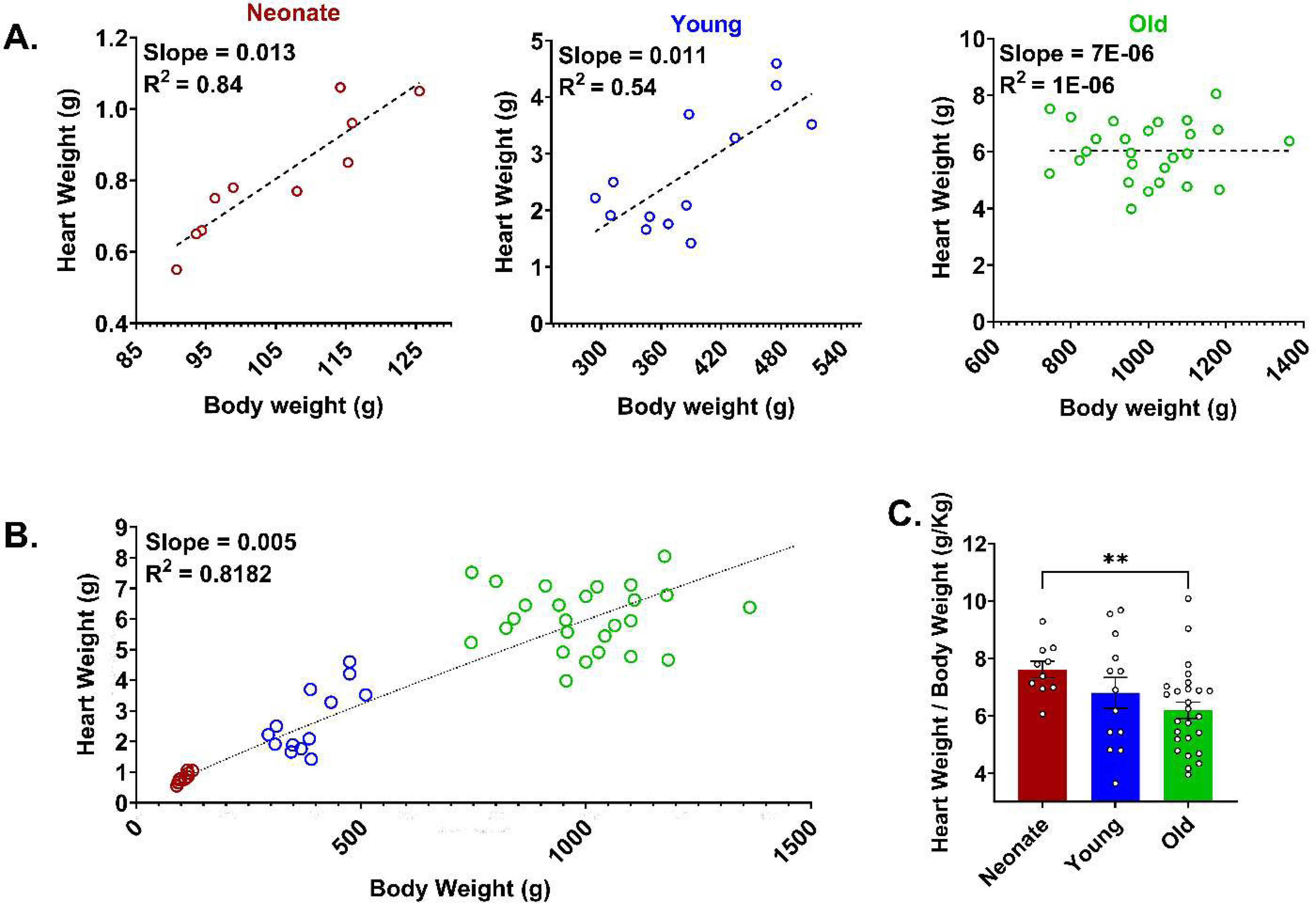
Heart weight-body weight relationship. **A)** The linear regression of normal heart weight versus body weight in neonatal (n=10), younger adult (n=13), and older adult guinea pigs (n=26). **B)** The regression slope of heart weight and body weight across different age groups. **C)** The heart weight-to-body weight ratio in each age group (mean ± SEM). *Comparisons by Welch’s ANOVA (unequal standard deviations between groups) with multiple comparisons test, **p<0.01*.

### Distinct *in vivo* electrical properties between neonatal, younger adult, and older adult hearts

Both human^5,34^ and guinea pig^24,25^ studies have reported an age-dependent effect on ECG parameters. Although, to date, preclinical studies have largely been limited to adults – with no comparison between pediatric and adult guinea pigs. In this study, we analyzed *in vivo* ECG data from 32 anesthetized guinea pigs (10 neonates, 13 younger adults, 9 older adults). A typical neonatal ECG trace (lead I) exhibited fusion of the T-wave with the P-wave of the next beat. The T-wave was less pronounced in younger adult animals, but both the P-wave and T-wave were clearly distinguishable in older adult animals (**Figure 2A**). Collectively, a number of ECG metrics were found to be statistically significant between the neonatal, younger adult, and older adult age groups (**Figure 2B**). For example, the intrinsic heart rate of neonates (239.6±15.0 bpm) and younger adults (247.4±17.6 bpm) was significantly faster than older adults (200.7±29.1 bpm, p<0.005; see **Table 1**). Atrial activation time, indicated by the P duration, was significantly shorter in neonates compared to their adult counterparts (*neonate:* 15.6±3.3 ms; *younger adult*: 22.7±3.0 ms; *older adult*: 24.1±2.90 ms; p<0.001). Ventricular activation time, indicated by the QRS interval, was also shorter in neonates (49.2±10.3 ms) compared to the younger (63.5±13.3 ms, p<0.05) and older adults (77.8±13.5 ms; p<0.001). However, no significant difference in QTc was observed across the age groups, suggesting that repolarization time was similar.

**Figure 2.**
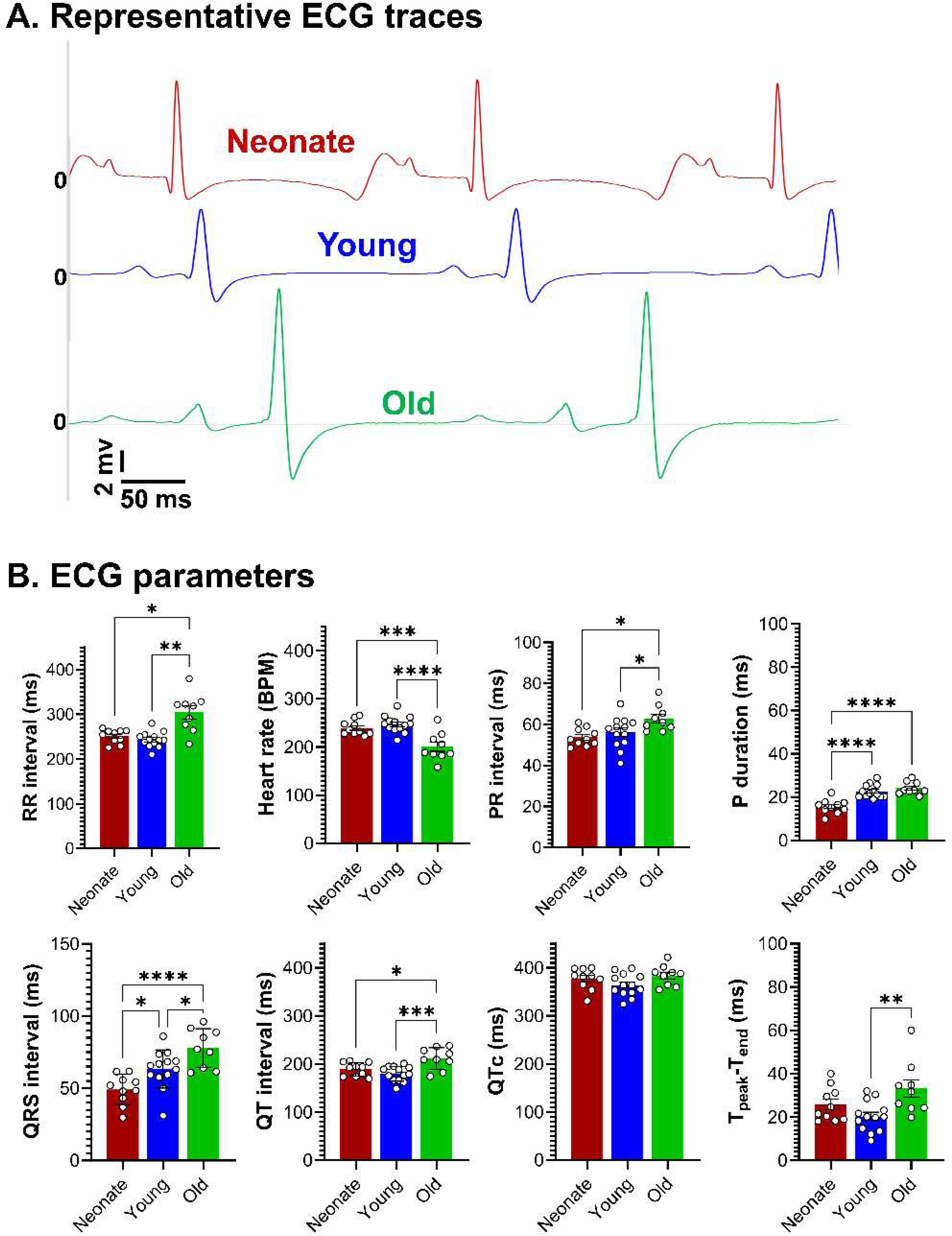
*In vivo* electrocardiogram (ECG) recordings across different age groups. **A)** Representative ECG traces (lead I) from neonatal (red), younger adult (blue), and older adult guinea pigs (green). ECGs were recorded in anesthetized animals using subcutaneous needle electrodes. **B)** Comparison of ECG metrics between neonates (n=10), young adults (n=13), and older adults (n=9). *All comparisons by ordinary one-way ANOVA with multiple comparisons test (with exception of Welch’s ANOVA for RR interval due to unequal standard deviations). Individual replicates shown; values reported as mean ± SEM. *p<0.05, **p<0.01, ***p<0.005, ****p<0.001*

**Table 1.**
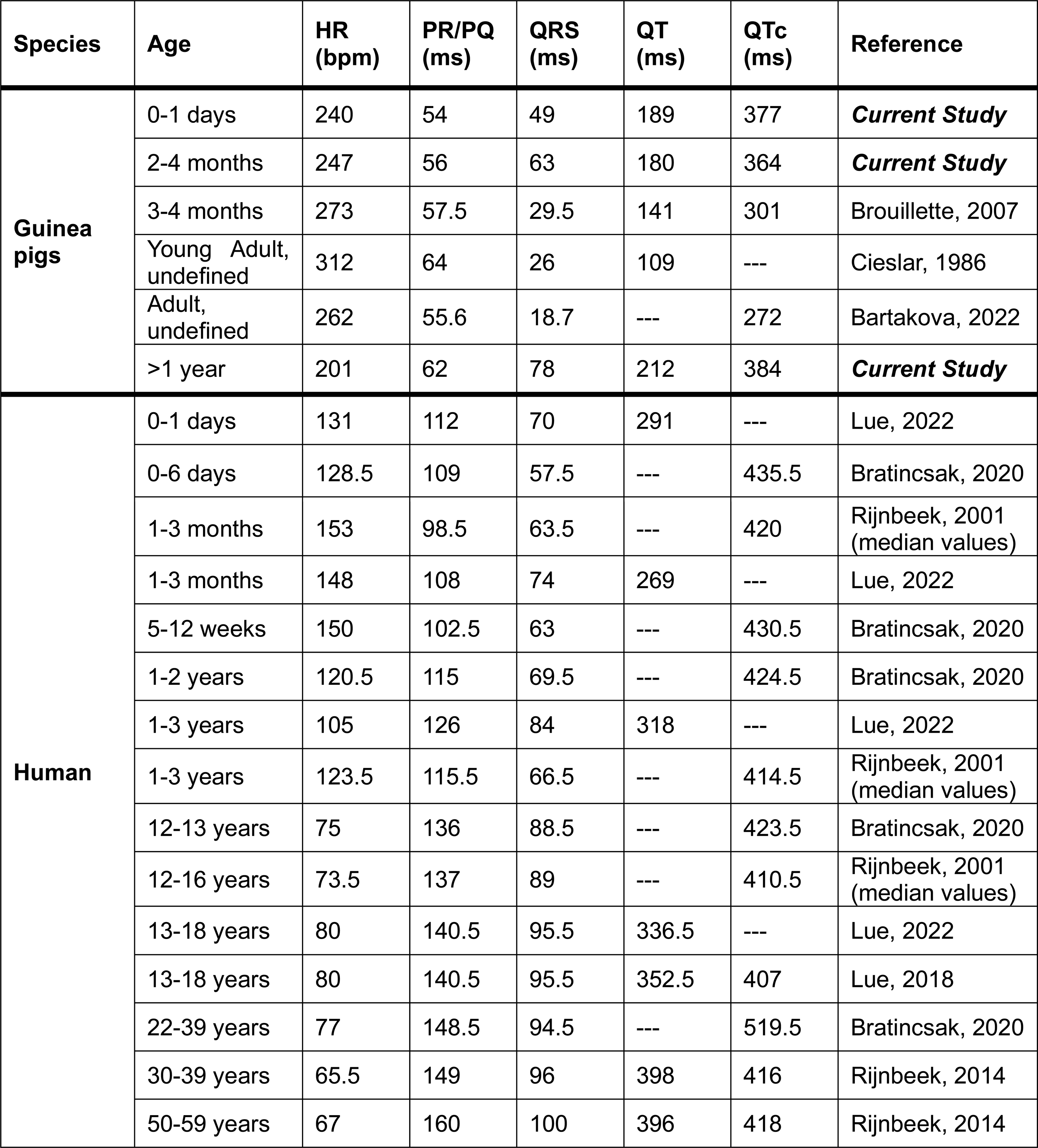
Mean values for electrocardiogram parameters in guinea pigs and humans. Male and female measurements were averaged, where applicable. Note that different measurement metric definitions and correction formulas between studies may account for variations in QRS, QT, and QTc values.

| Species | Age | HR (bpm) | PR/PQ (ms) | QRS (ms) | QT (ms) | QTc (ms) | Reference |
| --- | --- | --- | --- | --- | --- | --- | --- |
| Guinea pigs | 0-1 days | 240 | 54 | 49 | 189 | 377 | *Current Study* |
|  | 2-4 months | 247 | 56 | 63 | 180 | 364 | *Current Study* |
|  | 3-4 months | 273 | 57.5 | 29.5 | 141 | 301 | Brouillette, 2007 |
|  | Young Adult, undefined | 312 | 64 | 26 | 109 | --- | Cieslar, 1986 |
|  | Adult, undefined | 262 | 55.6 | 18.7 | --- | 272 | Bartakova, 2022 |
|  | >1 year | 201 | 62 | 78 | 212 | 384 | *Current Study* |
| Human | 0-1 days | 131 | 112 | 70 | 291 | --- | Lue, 2022 |
|  | 0-6 days | 128.5 | 109 | 57.5 | --- | 435.5 | Bratincsak, 2020 |
|  | 1-3 months | 153 | 98.5 | 63.5 | --- | 420 | Rijnbeek, 2001 (median values) |
|  | 1-3 months | 148 | 108 | 74 | 269 | --- | Lue, 2022 |
|  | 5-12 weeks | 150 | 102.5 | 63 | --- | 430.5 | Bratincsak, 2020 |
|  | 1-2 years | 120.5 | 115 | 69.5 | --- | 424.5 | Bratincsak, 2020 |
|  | 1-3 years | 105 | 126 | 84 | 318 | --- | Lue, 2022 |
|  | 1-3 years | 123.5 | 115.5 | 66.5 | --- | 414.5 | Rijnbeek, 2001 (median values) |
|  | 12-13 years | 75 | 136 | 88.5 | --- | 423.5 | Bratincsak, 2020 |
|  | 12-16 years | 73.5 | 137 | 89 | --- | 410.5 | Rijnbeek, 2001 (median values) |
|  | 13-18 years | 80 | 140.5 | 95.5 | 336.5 | --- | Lue, 2022 |
|  | 13-18 years | 80 | 140.5 | 95.5 | 352.5 | 407 | Lue, 2018 |
|  | 22-39 years | 77 | 148.5 | 94.5 | --- | 519.5 | Bratincsak, 2020 |
|  | 30-39 years | 65.5 | 149 | 96 | 398 | 416 | Rijnbeek, 2014 |
|  | 50-59 years | 67 | 160 | 100 | 396 | 418 | Rijnbeek, 2014 |

### Ventricular activation pattern and conduction velocity differs between neonates and adults

In humans, in utero development and postnatal cardiac maturation is marked by increased expression and alignment of ventricular gap junction proteins (connexin 40, 43)^35–37^. Gap junction remodeling has also been observed across developmental stages in canines, with connexin protein expression localized to the intercalated discs of adult ventricular cardiomyocytes – coinciding with a larger sodium current density (I_Na_) as compared to neonates^38,39^. This is an important developmental adaptation, as changes in connexin expression and I_Na_ density impact both the directionality and speed of electrical propagation by altering cell-to-cell electrical coupling^40^. Accordingly, we investigated ventricular activation patterns and measured CV in neonatal and adult guinea pigs. When pacing was applied to the center of the epicardium, neonatal hearts displayed a circular activation pattern while adult hearts had an elliptical activation pattern (**Figure 3A**). This activation pattern may be attributed to a disperse connexin-43 expression pattern in the neonatal myocardium, as compared with adults (**Figure 3D**). We also measured the maximal apex-to-base CV in response to dynamic pacing (S1-S1; **Figure 3B**). Age-dependent differences in CV were observed at all tested pacing frequencies (220-120 ms PCL; **Figure 3C**). Neonatal hearts consistently had the slowest CV measurements, while the fastest CV measurements were observed in younger adults. For example, at 200 ms PCL, neonatal CV measured 40.6±4.0 cm/s, which was slower than both younger adults (61.6±9.3 cm/s, p<0.001) and older adults (53.6±9.2 cm/s, p<0.005).

**Figure 3.**
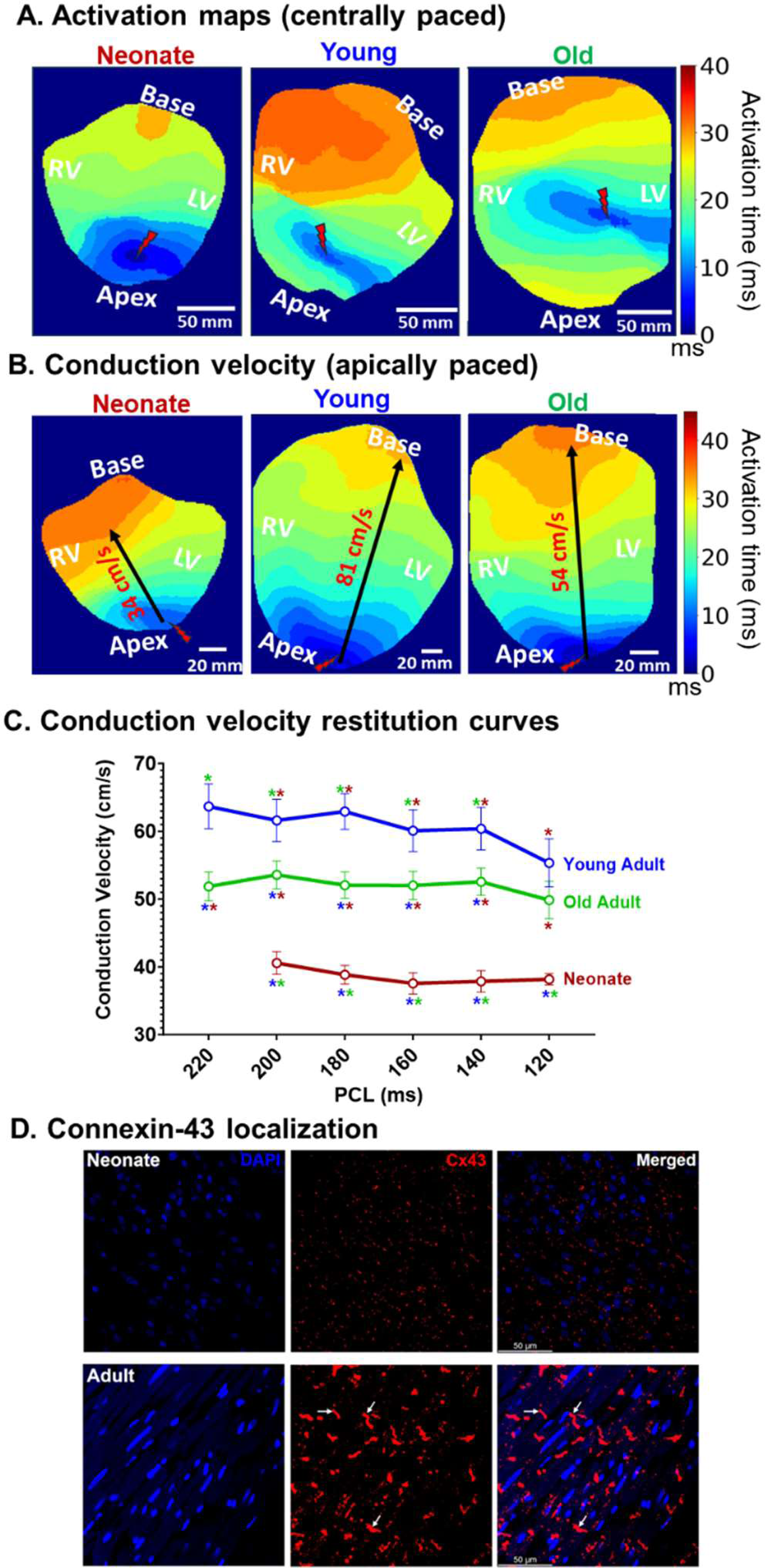
Ventricular activation dynamics differ across guinea pig age groups. **A)** Illustrative examples of ventricular AP activation maps (time of dV/dt_max_) in a neonate, younger adult, and older adult guinea pig heart. Electrical pacing (160 ms PCL) was applied near the geometric center, revealing a distinct activation pattern in neonatal hearts. **B)** Illustrative examples of epicardial CV measured from the earliest to latest activation site, as indicated by the single CV vector (black arrow). Electrical pacing (160 ms PCL) was applied near the apex. **C)** Epicardial CV restitution curves during S1-S1 pacing (220 to 120 ms PCL), featuring neonates (red, n=7), younger adults (blue, n=11), and older adults (green, n=21). **D)** Illustration of connexin-43 (red) localization in ventricular myocardium from neonatal and adult guinea pig; nuclei stained with DAPI (blue). 50 μm scale. *Replicate values reported as mean ± SEM. Comparisons by two-way ANOVA with multiple comparisons; *p<0.05 compared to neonate (red), young adult (blue), or old adult (green)*.

### Cardiac electrical restitution properties are modified by age

Restitution properties detail the rate-dependent adaptations in the cardiac AP and/or CaT, which are important determinants of arrhythmia susceptibility^31,41^. Previous studies have reported AP and CaT restitution in the guinea pig heart in response to both dynamic (S1-S1) and extrasystolic (S1-S2) stimulation^23,42–44^, although age-dependent variations have yet to be investigated. Collectively, our results show that the epicardial APD and CaD lengthens with increasing age and that this trend is most evident at slower pacing frequencies (**Figure 4A, B**). For example, at a slower pacing rate (200 ms PCL), neonatal hearts had a significantly shorter APD50 (86.0±3.9 ms) and CaD50 (94.3±3.3 ms) compared to both younger (*APD50:* 96.8±5.4 ms, p<0.05; *CaD50:* 99.4±2.4 ms, <0.005) and older adults (*APD50:* 107.7±2.0 ms, p<0.05; *CaD50*: 109.4±1.4 ms, p<0.001). Such age-dependent differences were smaller in magnitude at faster pacing frequencies. At a slightly faster rate (160 ms PCL), neonatal heart measurements (*APD50:* 81.0±3.4 ms, *CaD50:* 86.1±2.6 ms) were comparable to younger adult hearts (*APD50:* 88.7±5.4 ms, *CaD50:* 82.8±1.8 ms), but remained significantly different from older adult hearts (*APD50*: 96.5±1.8 ms, p<0.001; *CaD50:* 89.2±2.0 ms, p<0.001).

**Figure 4.**
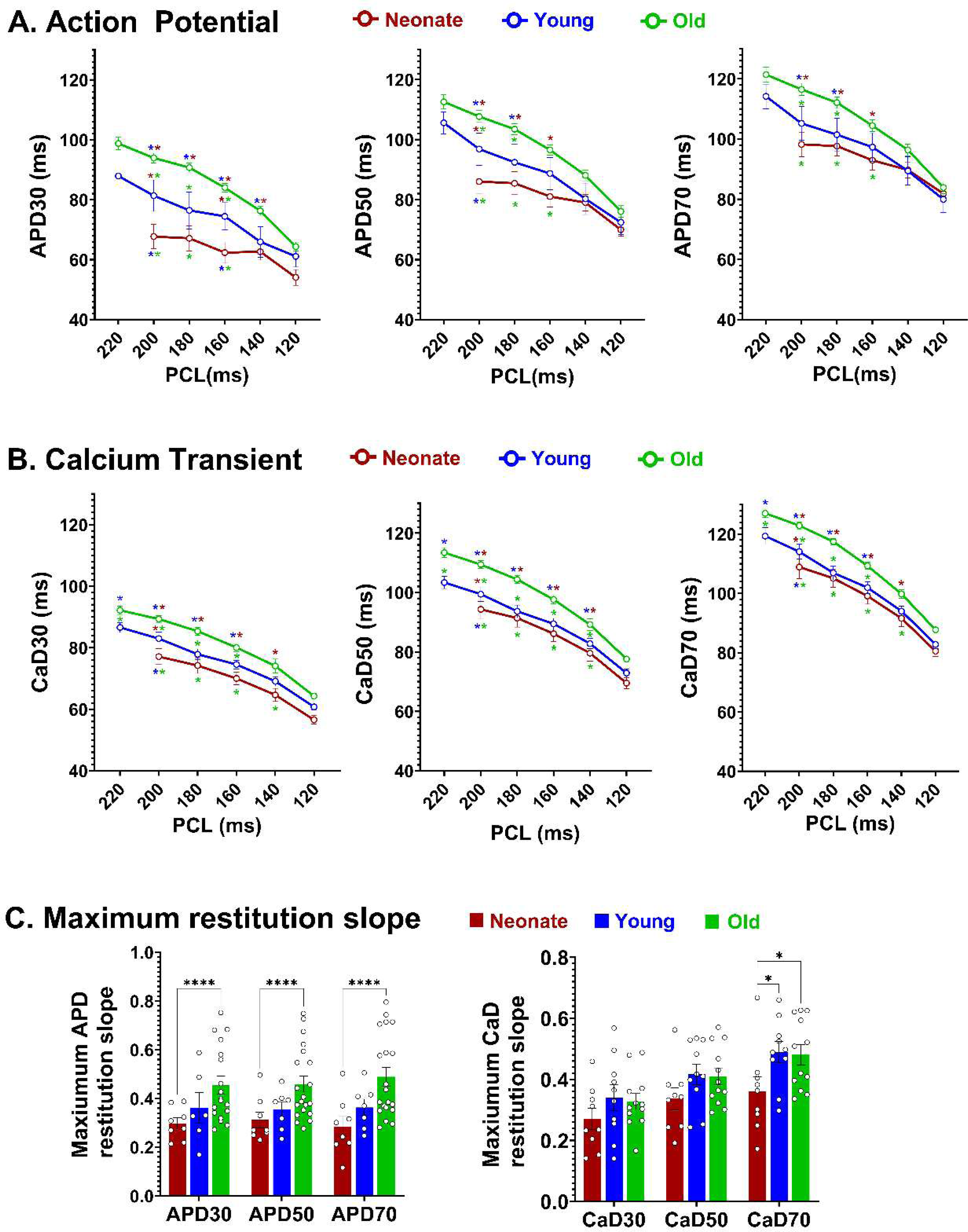
Cardiac electrical restitution properties in response to dynamic stimulation. **A)** Action potential duration restitution curves at APD30, APD50, APD70 in neonatal (n=8), younger adult (n=6-7), and older adult hearts (n=18-21). **B)** Calcium transient duration restitution curves at CaD30, CaD50, CaD70 in neonatal (n=9), younger adult (n=10), and older adult hearts (n=12). **C)** Maximum restitution slopes; individual replicates shown. Ventricles were dynamically paced (S1-S1) at the apex; the pacing cycle length was decremented from 220 to 120 ms at 20 ms intervals. *Values reported as mean ± SEM. Comparisons by two-way ANOVA with multiple comparisons; *p<0.05, ****p<0.001 compared to neonate (red), young adult (blue), or old adult (green)*.

Similarly, the maximum slope of the each APD restitution curve (APD30, APD50, APD70) at S1-S1 pacing exhibited significant age-dependent differences between neonates and older adults, but not younger adults (**Figure 4C**). For example, the maximum APD70 restitution slope in neonates measured (0.29±0.04, n=8), which was significantly smaller than in older adult hearts (0.49±0.04, n=20, p<0.0001) but not different from younger hearts (0.36±0.03, n=7). In contrast, only the CaD70 restitution curve differed between age groups. Neonates had a significantly flatter restitution curve (0.36±0.05, n=9) compared to both younger (0.49±0.04, n=10, p<0.05) and older adults (0.48±0.03, n=12, p<0.05). The maximum restitution slope for CaD30 and CaD50 trended flatter for neonatal hearts compared with adults, but the difference was not statistically significant (**Figure 4C**).

### Neonatal epicardial repolarization displays both regional and temporal variations with dynamic stimulation

In humans, ECG recordings have revealed variations in cardiac electrical properties that coincide with postnatal heart development – including an increase in heart rate (131±12.8 to 148±15.6 bpm), shortening of the QT interval (291±26.6 to 259±17.9 ms), and change in the frontal plane QRS axis (135 to 110°) between day 0 and day 30 of the neonatal period ^5,45^. Similar developmental shifts in ECG and APD parameters have been reported in postnatal rats (day 0 to day 14)^20^. Accordingly, we investigated whether such dynamic changes in cardiac electrical properties occurred in the neonatal guinea pig. Collectively, we found that postnatal heart development was characterized by a shorter APD, which was most evident at slower pacing rates (S1-S1: 200-140 ms PCL; **Figure 5A**). For example, from postnatal day 0 to day 1 the APD30 shortened from 73.4±5.5 to 60.0±2.5 ms (p<0.05) and APD70 shortened from 104.3±3.4 to 90.0±5.8 ms (p<0.05) at 200 ms PCL. However, APD values were comparable between postnatal day 0 and day 1 hearts at faster pacing rates (120 ms PCL). There was no significant difference in CaD measurements (CaD30, 50, 70) between postnatal day 0 and day 1 hearts – although CaD70 trended longer in day 0 hearts at the slower pacing rate (200 ms PCL; **Figure 5B**).

**Figure 5.**
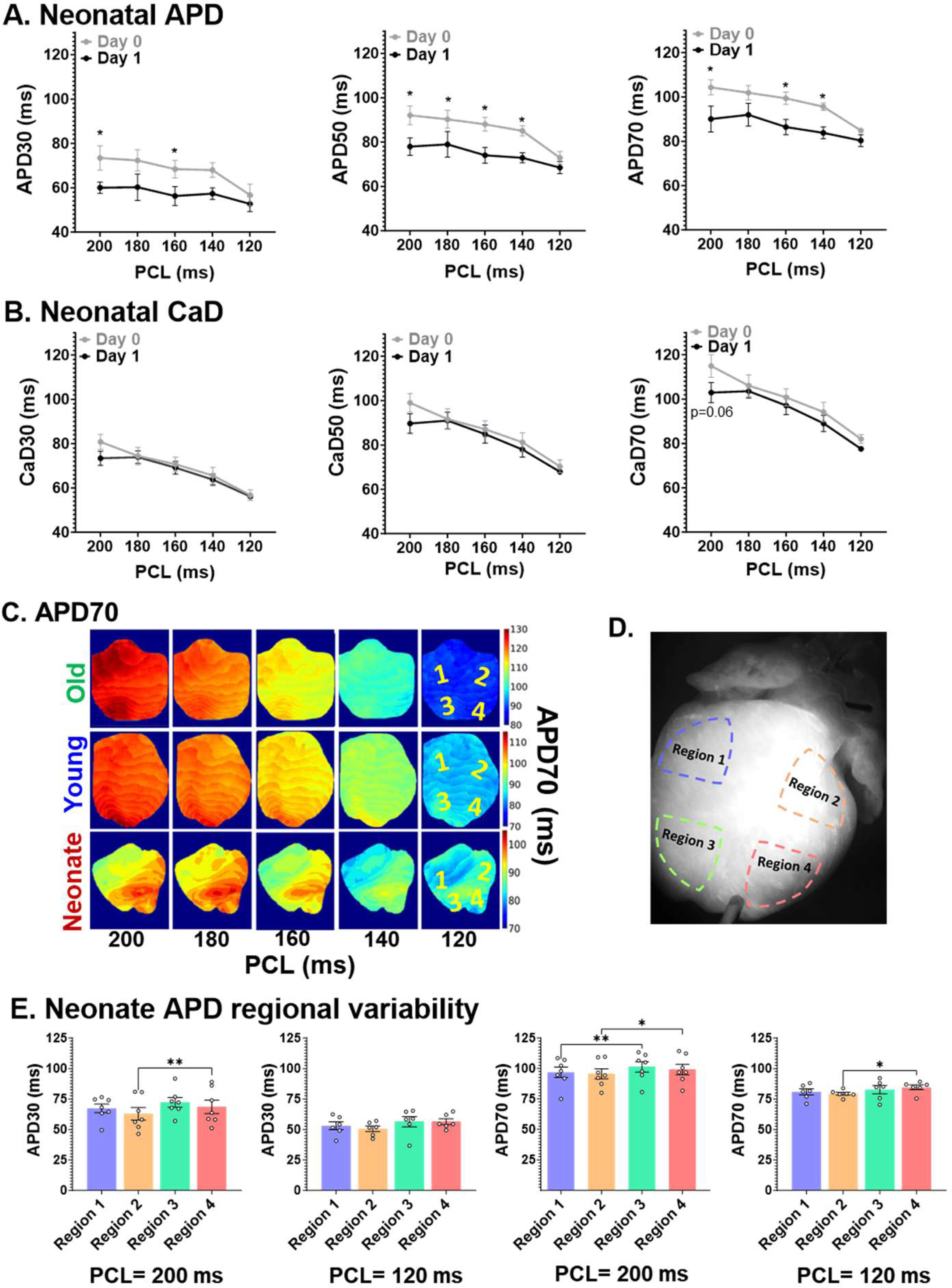
Intragroup and regional variability in neonatal guinea pig hearts (dynamic stimulation). **A)** Action potential duration (APD) and **B)** calcium transient duration restitution curves generated from neonatal hearts isolated on postnatal day 0 (n=4-5) and postnatal day 1 (n=4). Dynamic pacing (S1-S1) was applied to the apex; the pacing cycle length (PCL) was decremented from 200 to 120 ms. *Values reported as mean ± SEM. Comparison by two-way ANOVA with multiple comparisons, *p<0.05.* **C)** Illustrative APD70 maps of a neonate, younger adult, and older adult heart in response to dynamic S1-S1 apical pacing. Note the regional variability in the neonatal heart. **D)** Schematic of four epicardial ventricular regions of interest. **E)** APD30 and APD70 measurements from neonatal hearts, collected in response to a slow pacing rate (200 ms PCL, n=7) and faster pacing rate (120 ms PCL, n=6). *Values reported as mean ± SEM. Comparison by one-way (matched) ANOVA with multiple comparisons; *p<0.05, **p<0.01 between different regions*.

A previous study documented transmural and apicobasal APD heterogeneity in tissues isolated from adult guinea pig hearts^46^. Accordingly, we performed regional analysis of repolarization maps and noted an intriguing aspect of neonatal epicardial electrophysiology. As illustrated in **Figure 5C**, neonatal APD70 maps displayed regional heterogeneity at multiple different pacing rates – with longer APDs typically measured near the apex (region of interest 3 & 4; **Figure 5D**). For example, at a slow pacing rate (200 PCL), APD30 and APD70 were slightly longer in Region 4 (left apex; *APD30:* 68.6±5.6 ms, *APD70*: 99.2±11.4 ms) as compared to Region 2 (left basal; *APD30:* 62.8±3.6, *APD70:* 95.6±10.8 ms; **Figure 5E**). In comparison, regional APD heterogeneity was not observed in either younger or older adults (**Figure 5C**, *compiled data not shown*). Additionally, regional CaD heterogeneity was not observed in any age group (*data not shown*).

### Age-dependent variations in cardiac electrical restitution properties are evident with extrasystolic stimulation

Distinct electrical restitution properties have been observed in adult guinea pigs in response to dynamic versus extrasystolic pacing, as demonstrated by APD restitution curves^44^. Accordingly, we investigated whether neonatal guinea pigs exhibit unique APD and/or CaD restitution characteristics with extrasystolic pacing. Notably, S1-S2 pacing captured at shorter intervals in neonatal hearts compared to adult hearts (**Figure 6A,B**). In neonatal hearts, the longest achievable S1-S2 interval was 150-140 ms, which increased to 200-180 ms in younger adults and 200-190 ms in older adults. Interestingly, the measured APD values in young and old adult hearts were nearly identical (**Figure 6A**) – yet, significant differences in CaD values were observed between these cohorts (**Figure 6B**). As an example, in adults the APD70 values were comparable at 200-180 ms pacing rate (*Young adult:* 108.5±2.8, *Old adult:* 112.8±2.9 ms), but young adults had slightly shorter CaD70 values (113.2±2.6 ms) compared with old adults (123.7±3.3 ms, p<0.05). Next, we calculated the maximum APD and CaD restitution slopes in response to extrasystolic pacing. Collectively, neonatal hearts exhibited flatter APD and CaD restitution slopes compared to adults (**Figure 6C**). For instance, the maximum APD70 and CaD70 restitution slopes of neonates (*APD70:* 0.54±0.09 ms, n=6, *CaD70:* 0.51±0.04 ms, n=7) was significantly smaller than the adults (*Old APD70:* 0.95±0.7 ms, n=10, p<0.0001; *Young APD70*: 0.85±0.04 ms, n=5, p<0.05; *Old CaD70:* 1.18±0.14, n=7, p<0.0001; *Young CaD70:* 0.89±0.11, n=5, p<0.05).

**Figure 6.**
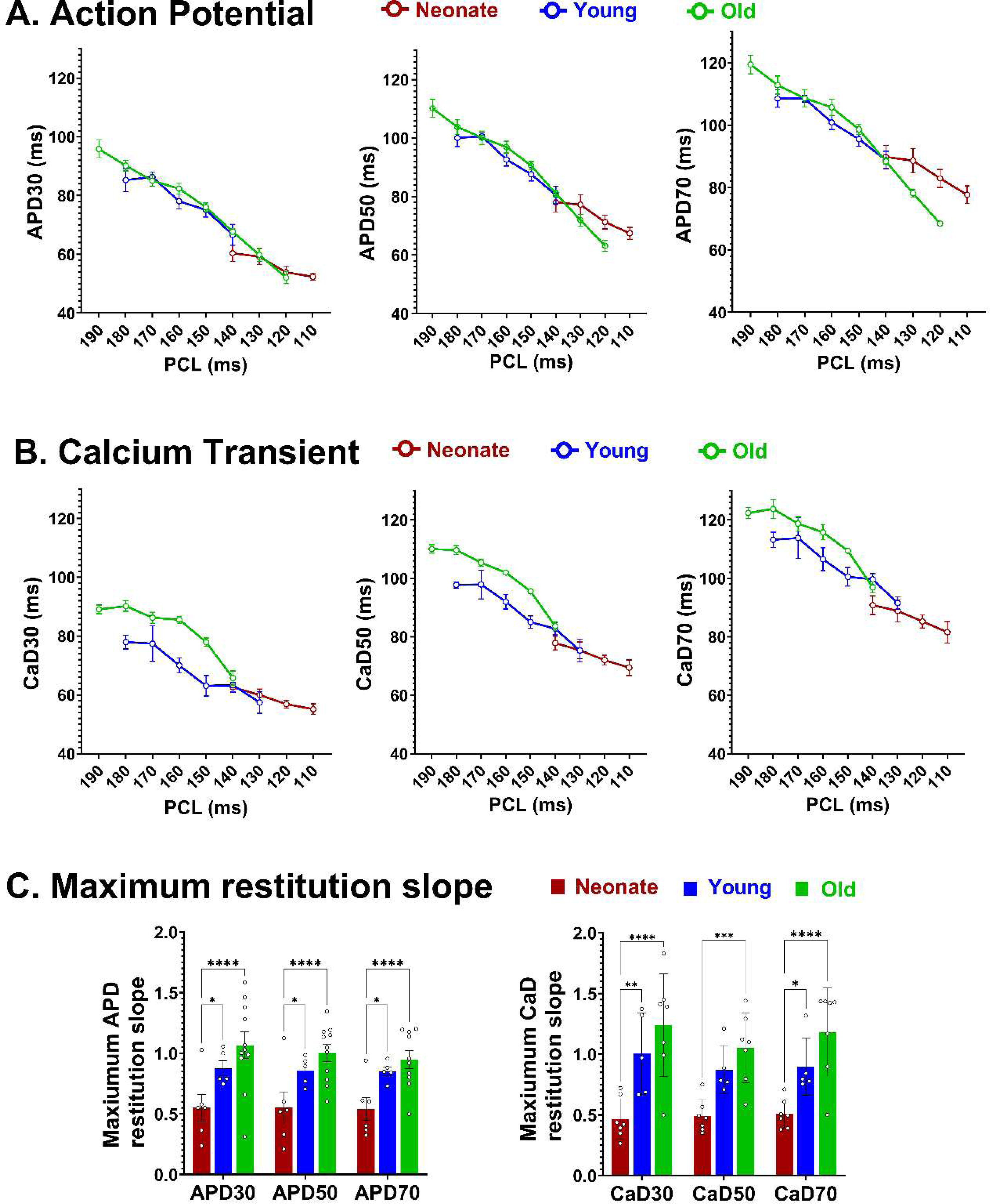
Cardiac electrical restitution properties in response to extrasystolic stimulation. **A)** Action potential duration restitution curves at APD30, APD50, APD70 in neonatal (n=5-6), younger adult (n=5), and older adult hearts (n=6-11). **B)** Calcium transient duration restitution curves at CaD30, CaD50, CaD70 in neonatal (n=5-6), younger adult (n=5), and older adult hearts (n=6). **C)** Maximum restitution slopes; individual replicates shown. Extrasystolic stimulation was applied to the ventricle (S1-S2) at the apex. Note the different pacing cycle length ranges that were achievable in neonatal versus adult hearts. *Values reported as mean ± SEM. Statistical comparisons between slope measurements, as determined by one-way ANOVA with multiple comparisons; *p<0.05, **p<0.01*.

Since dynamic stimulation (S1-S1) produced regional heterogeneity in epicardial APs in neonatal hearts, we assessed whether this heterogeneity persisted with extrasystolic stimulation (S1-S2). As shown in **Figure 7A,B**, neonatal epicardial APD50 maps (S2 beat shown) and traces suggested longer APDs in the right ventricle compared to the left ventricle. This observation was supported by regional APD restitution curves (**Figure 7C**). For example, at a slower S1-S2 pacing interval (150-140 ms), APD70 was shorter in the left ventricular base (*Region 2:* 84.9±2.2 ms) and left ventricular apex (*Region 4:* 88.7±2.7 ms) – as compared to the right ventricular base (*Region 1*: 90.9±5 ms) and right ventricular apex (*Region 3:* 94.7±5 ms). Regional heterogeneity was much less pronounced in neonatal heart CaD measurements, with only slightly shorter CaD70 values detected in the left ventricular base (*Region 2)*. In contrast to neonates, younger adult hearts did not exhibit any significant regional heterogeneity in either APD or CaD (**Figure 7D**) – while older hearts displayed only minor regional variability in APD70 at a slower S1-S2 pacing interval (**Figure 7E**).

**Figure 7.**
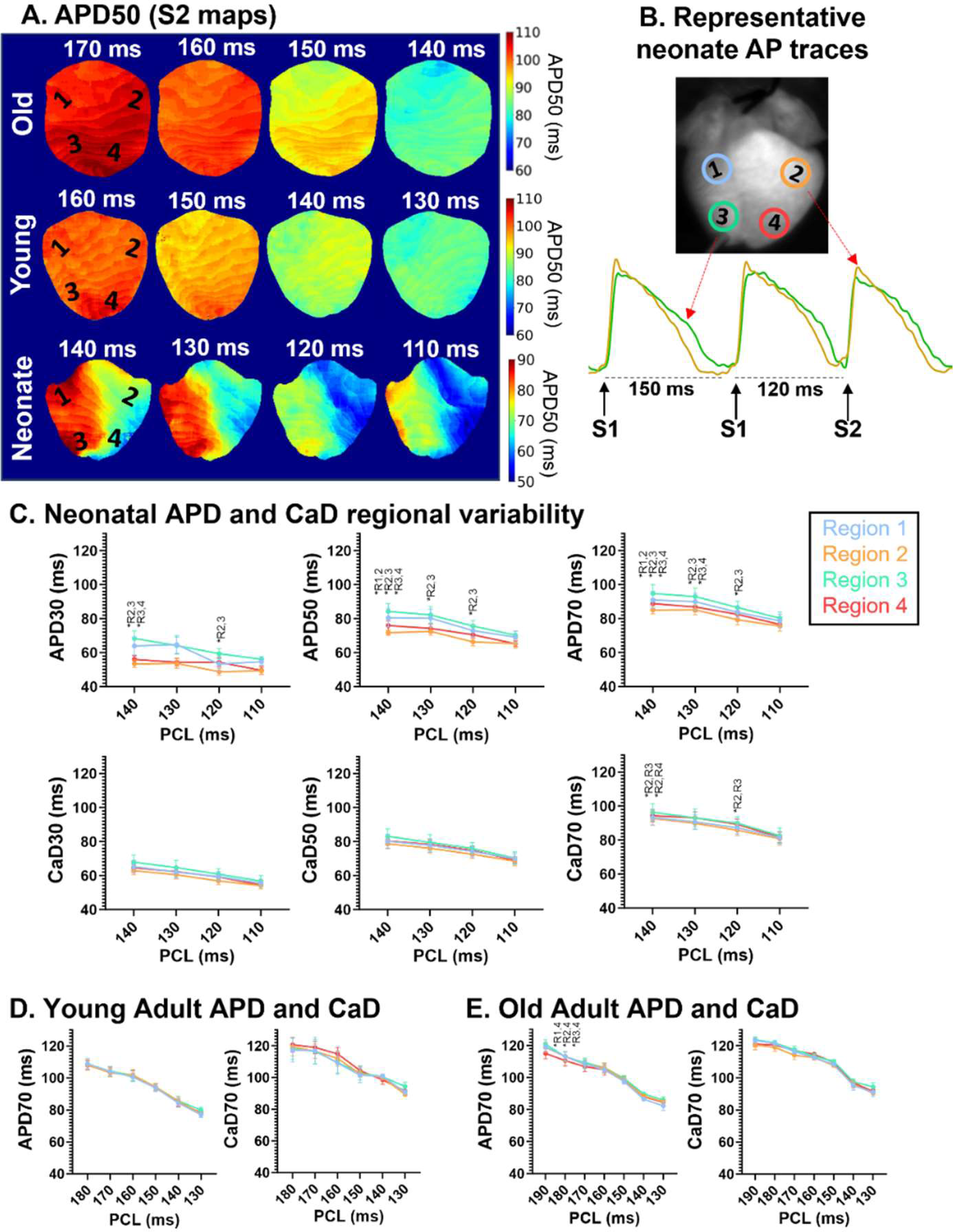
Regional variability in neonatal guinea pig hearts (extrasystolic stimulation). **A)** Illustrative APD50 maps of a neonate, younger adult, and older adult heart in response to extrasystolic S1-S2 apical pacing (maps highlight the “S2” beat). **B)** Representative AP traces from two regions of the neonatal heart (Region 2: left basal in orange vs Region 3: right apex in green). **C)** Regional heterogeneity in action potential duration (APD) and calcium transient duration (CaD) restitution curves generated from neonatal hearts (n=6-7). **D, E)** APD70 and CaD70 restitution curves generated from younger (n=6-9) and older adult hearts (n=6-8). *Values reported as mean ± SEM. Comparison by two-way ANOVA with multiple comparisons, *p<0.05 between denoted specific regions*.

### Distinct excitation-contraction coupling properties in neonatal hearts

Excitation-contraction coupling (ECC) refers to a series of signaling processes whereby an AP triggers a rise in intracellular calcium, allowing calcium ions to interact with contractile protein machinery to initiate muscle contraction^47^. ECC has been studied in isolated adult guinea pig cardiomyocytes through both experimental^48,49^ and computational approaches^50^. However, to date, there is limited information on ECC in the context of the intact heart. Using our previously described methodology^26^, we generated action potential-calcium transient (AP-CaT) latency maps using optical signals collected from guinea pig hearts. As shown in **Figure 8A**, latency maps from adult hearts reveal negative coupling (APD30 > CaD30) during early repolarization and positive coupling (APD70 < CaD70) during late repolarization. In contrast, in the neonatal ventricular epicardium, AP-CaT coupling is consistently positive with CaD > APD at all repolarization phases. This observation is further substantiated by quantitative measurements of mean latency plots at variable S1-S1 pacing rates (**Figure 8B**). As the pacing cycle interval is decremented (200 to 120 ms), AP-CaT latency remains positive in neonatal hearts across different repolarization phases (n=4). While in younger (n=5) and older adult hearts (n=8), CaD30-APD30 is negative for most pacing rates, close to zero for CaD50-APD50, and shifts to a more positive coupling for CaD70-APD70. These results suggest a distinct ECC mechanism that is unique to neonatal ventricles.

**Figure 8.**
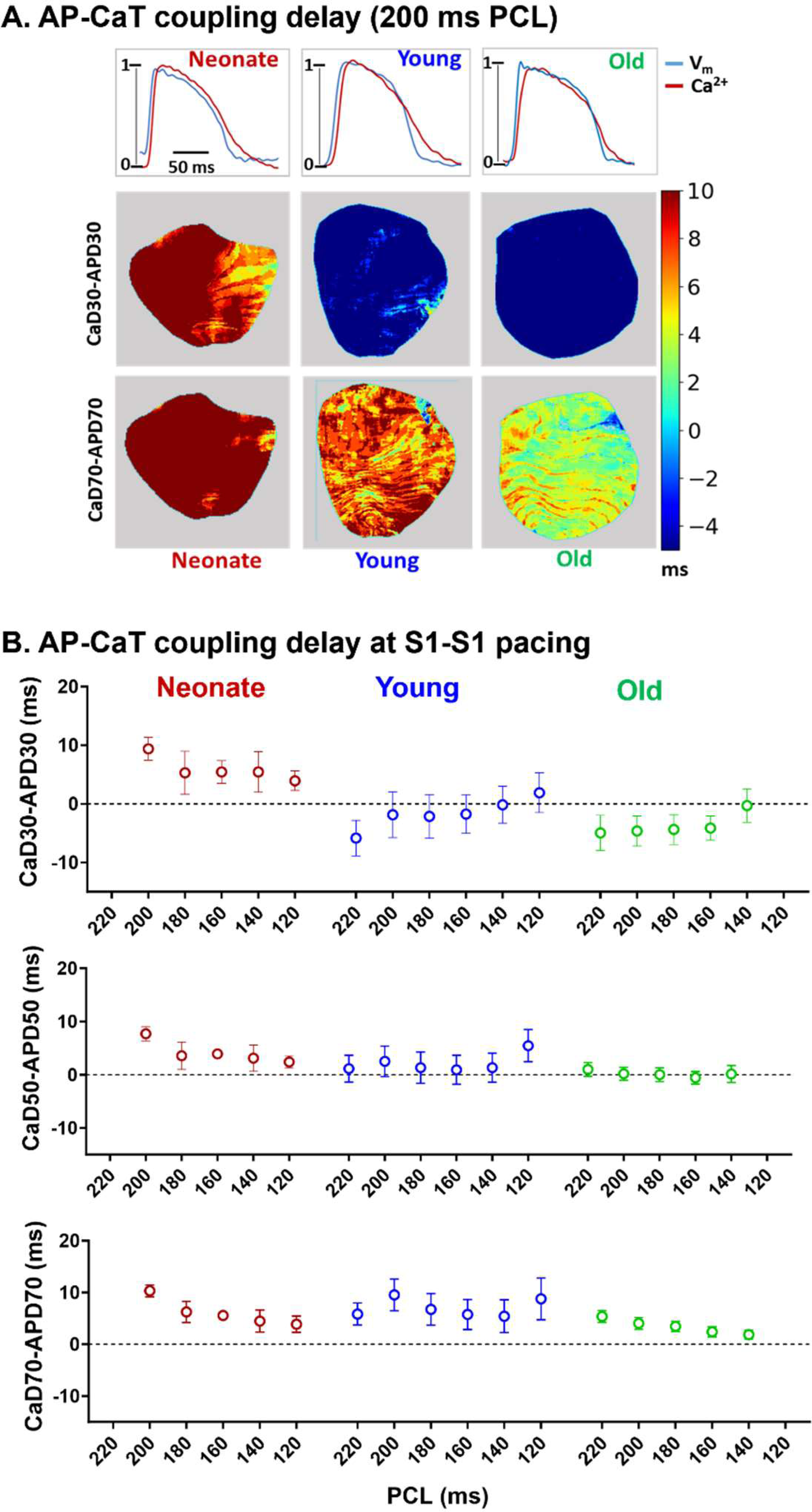
Action potential and calcium transient coupling characteristics across guinea pig age groups. **A)** Top: Representative traces of AP and CaT recorded simultaneously from a neonate, younger adult, and older adult heart. Bottom: Representative AP-CaT latency maps (CaD-APD) measured at 30% and 70% duration. **B)** Quantitative comparison of AP-CaT latency at 30%, 50%, and 70% duration in neonatal (n=4), younger adult (n=6), and older adult (n=8) guinea pigs. Positive values indicate that the CaD is longer, while negative values indicate that the APD is longer for a given measurement. All measurements recorded in response to dynamic (S1-S1) ventricular epicardial pacing at the apex. *Values reported as mean ± SEM*.

## Discussion

This ambispective cohort study aimed to explore whether the electroanatomical behavior of the heart differs between the early postnatal period and adulthood in guinea pigs. Collectively, our investigation involved measuring heart weight:body weight, recording *in vivo* electrocardiograms using subcutaneous electrodes, and performing optical mapping and electrophysiology studies *ex vivo* using Langendorff-perfused heart preparations. Our study unveiled unique electroanatomical characteristics in neonatal guinea pig hearts, and our main findings include: 1) neonatal hearts exhibit a more robust and steeper relationship between body weight and heart weight as compared to adults, 2) *in vivo* ECG metrics differ between age groups, 3) neonatal hearts display unique ventricular epicardial activation patterns as compared to adults, 4) neonatal hearts respond differently than adult hearts to dynamic and extrasystolic stimulation, and exhibit distinct AP and CaT restitution characteristics, 5) neonatal hearts display APD heterogeneity, but CaD is more homogeneous across the ventricular epicardium, 6) neonatal hearts have distinct ECC characteristics with CaD > APD at different phases of repolarization.

### Robust allometric relationship between heart weight and body weight in neonatal guinea pigs

The allometric relationship between biological variables and body mass has been studied across multiple species^51,52^. This relationship is described by an exponential fit, with the exponent commonly found to be less than 1 or close to 3/4^51–54^. Cardiovascular variables have also been shown to scale with body mass in mammals^53^. Specifically, an allometric relationship between body mass and cardiac electrophysiology has been demonstrated using ECG parameters across different species^55^. In the current study, a similar allometric correlation was found between guinea pig heart weight and body weight, with an exponential fit (R^2^ = 0.82) using all heart and body weight measurements across the different age groups (**Figure 1**). Interestingly, the power exponent was determined to be 0.89, which is close to the allometric exponent value reported for humans (0.85)^56^. This similarity further supports the robustness of the guinea pig model for studying cardiovascular development and maturation, as highlighted by others^21,22^. When split into respective age groups, a linear fit revealed a strong correlation (0.84) and a steeper slope (0.013) in neonates, as compared to adults (young: 0.011, old: 7E-06). Neonates also presented with a larger heart-to-body weight ratio as compared to older adults. These results indicate that heart mass does not increase with additional body weight in older animals. In older adult guinea pigs, the absence of allometry is likely due to increasing body weight as a function of body fat deposition^57^, which can induce significant inter-subject variability.

### Electrocardiographic parameters are age-dependent in guinea pigs

ECG recordings are frequently employed as an accurate and non-invasive method to characterize the electrical behavior of the entire heart, which provides invaluable diagnostic information in the clinical setting. Human ECG studies have reported age-dependent characteristics, which highlight the impact of heart muscle growth on electrophysiology (**Table 1**)^5,34,58–60^. Briefly, the human heart rate increases after birth within the first 30 days of life (0-1 day: 131 vs 30 days: 148 BPM), and subsequently slows throughout childhood (1-3 years: 105 BPM), adolescence (13-18 years: 80 BPM), and adulthood (30-39 years: 65.5 BPM)^5,34^. In the current study, we noted a similar trend in our experimental guinea pig model, wherein heart rate increased after birth in younger animals and then subsequently slowed in older adult animals (**Table 1**). Other age-specific ECG parameters that shifted with advanced age included an increase in the P duration (atrial conduction time), PR interval (atrioventricular conduction time), QRS duration (ventricular activation time; **Figure 2**) – which is consistent with human developmental trends. These developmental changes are likely explained by the increasing heart mass during the perinatal period and early adulthood, which requires more time for electrical signals to propagate. While in later adulthood, the deposition of cardiac adipose tissue^61^ and/or increased fibrosis^62^ can slow electrical propagation. It is important to note that variations in ECG recording protocols (e.g., invasive vs non-invasive approach, use/type of anesthetics, metric definitions, and correction formulas) can contribute to discrepancies in reported ECG measurements^24^. Nevertheless, our adult guinea pig ECG data falls within the range of previously reported values (**Table 1**)^63–65^. Further, the current study expands our current knowledge by incorporating a neonatal guinea pig cohort for ECG measurements, which has not previously been reported.

### Neonatal ventricular activation is characterized by a slow, circular conduction pattern

In the current study, we observed a circular conduction pattern in neonatal guinea pig hearts when external pacing was applied centrally to the ventricular epicardium. In contrast, adult hearts exhibited a prototypical elliptical activation pattern – which is linked to age-dependent gap junction remodeling that increases connexin expression at the cardiomyocyte termini^66^. In comparison, connexin proteins are spatially dispersed throughout the neonatal cardiomyocyte^66,67^, which likely explains the circular activation pattern observed in our study. Our immunostaining results serve as proof of concept, supporting the potential molecular mechanism underlying this unique activation pattern – with dispersed connexin-43 expression in neonatal ventricular tissue and more focused expression at the cardiomyocyte termini in adult tissue (**Figure 3D**).

This distinct epicardial activation pattern in neonates was also reflected in ventricular CV measurements, which was significantly slower as compared to adult counterparts (**Figure 3C**). Slower CV in neonates can be attributed to weak intercellular electrical coupling (connexin expression noted above), less sodium current^38^, and limited sodium channel localization at the intercalated discs^36^. At the other end of the age spectrum, older adult hearts display slowed CV (compared to younger adults) that may be attributed to adipose tissue deposition^61^ and the accumulation of fibrotic tissue^62^. Notably, our reported values for adult CV measurements and the relatively flat CV restitution curve are both in agreement with previous reports^43,68,69^.

### Neonatal hearts have shorter APDs, shorter CaDs, and flattened repolarization restitution curves

When dynamically paced at the ventricular apex, neonatal hearts had significantly shorter APDs and a flattened repolarization restitution curve compared to adult hearts (**Figure 4**). Our results align with previous studies that measured APDs from isolated cells, which reported shorter APDs in neonatal versus adult ventricular cardiomyocytes^70,71^. In our study, neonatal hearts also displayed shorter CaDs and a flattened restitution slope compared to adult hearts – particularly at CaD70 (**Figure 4**). These results align with previous findings in neonatal cardiomyocytes, including the lack of well-developed SR storage^72^ and intracellular calcium dynamics that are largely mediated by extracellular calcium^73^. Moreover, immature cardiomyocytes rely more heavily on the sodium-calcium exchanger for intracellular calcium handling, as compared to SR-calcium handling channels and transporters^74^. Of interest, we also reported that the maximum APD and CaD restitution curve slopes were comparable between neonates and younger adults – suggesting that age-related rate-adaptation occurs later in guinea pig heart development (e.g., compared to conduction velocity). Whether this trend is preserved in other animals or in humans will require further study.

### Neonatal hearts exhibit temporal and regional variability in action potentials, but not in calcium transients

An unexpected finding in our study was a significant age-dependent change in the repolarization characteristics of neonatal hearts (**Figure 5**). When dynamically paced at the ventricular apex, longer APDs were measured in neonatal hearts isolated on postnatal day 0 and shorter APDs were measured on postnatal day 1. While there is no available data on cardiac electrophysiology during postnatal guinea pig development, single-cell studies have reported longer APDs in fetal versus neonatal cardiomyocytes ^70,71^. In humans, the QT interval (another metric of ventricular repolarization) also shortens during the early postnatal period from day 0 to day 30^5^. However, calcium handling properties remained unchanged between postnatal day 0-1, as indicated by nearly identical CaD restitution curves (**Figure 5B**).

Another unique finding of our study was an increase in neonatal APD heterogeneity during both dynamic (**Figure 5**) and extrasystolic (**Figure 7**) ventricular pacing. Epicardial APD heterogeneity was also more prominent during slower versus faster pacing rates. Prior studies have reported regional heterogeneity between the left versus right ventricle^75^ using isolated ventricular cardiomyocytes, and/or base versus apex^46^ in intact heart preparations from young guinea pigs. Although, in our study, regional heterogeneity was largely absent in adult animals (**Figure 5C**, **Figure 7D-E**). Discrepancies between studies could be related to the model system (isolated cardiomyocytes vs intact heart preparations), the recording technique (monophasic AP recording^46^ vs optical mapping), or the pacing protocol (e.g., pacing rate, dynamic vs extrasystolic). In our study, the S1-S1 pacing protocol resulted in apicobasal heterogeneity (**Figure 5E**), while the S1-S2 pacing protocol yielded more heterogeneity between the left and right ventricle (**Figure 7C**). Although transmural heterogeneity in repolarizing currents has been reported in adult humans^76^, a comparable study has not been performed on the neonatal guinea pig heart. Notably, we did not observe the same degree of spatial heterogeneity in CaDs recorded from neonatal guinea pig hearts.

### Neonatal excitation-contraction coupling is unidirectional while adults exhibit bidirectional coupling

The AP-CaT time latency reflects underlying ECC characteristics. In our study, the time latency (measured as CaD-APD at a specific repolarization phase) was consistently positive in neonatal hearts at all pacing cycle lengths (**Figure 8**). In contrast, adult guinea pig hearts had variable time latency patterns. At 30% and 50% repolarization, the time latency was often negative or nearly zero (indicating a longer AP with a comparatively shorter CaT). At 70% repolarization, the time latency became positive (indicating a longer CaT with comparatively shorter AP). This result suggests that unidirectional AP-CaT coupling is unique to neonatal guinea pig hearts, while in adults, the coupling is bidirectional. It has been reported that neonatal hearts lack a t-tubular system and well-developed SR storage ^72^, which indicates that intracellular calcium dynamics in this age group are primarily influenced by transmembrane voltage-dependent calcium channels (L-type/T-type) and sodium-calcium exchanger^74^. Accordingly, the intracellular calcium concentration is largely influenced by the extracellular calcium concentration^73^. Elevated extracellular calcium levels can ensure uniform calcium entry during systole, contributing to the observed unidirectional coupling.

### Limitations

The scope of our study was limited to developmental adaptations in cardiac electrophysiology and calcium handling between neonatal (postnatal day 0-1) and adult guinea pig hearts. As such, additional work is required to further investigate developmental trends that occur later in postnatal development. In vivo ECG recordings were performed under anesthesia; therefore, ECG measurements will be different from those recording in awake animals^24,77^. Further, our ECG measurements could be influenced by slight differences in the anesthetic response between individual animals and/or age groups. The bulk of our experiments were performed using excised, Langendorff-perfused heart preparations, which could conceal other age-dependent differences in cardiovascular physiology and/or autonomic regulation. Further, our ex vivo optical data sets were collected from the anterior ventricular epicardial surface, and as such, results may differ in other heart locations (e.g., posterior, endocardium, atrium). The location of the pacing electrode can influence activation patterns, and consequently, CV measurements. We attempted to consistently place the pacing electrode on the apex for AP and CaT measurements, and also recorded a second set of images with the electrode paced centrally to view activation patterns. However, we should note that exact placement on the smaller neonatal guinea pig heart posed some minor technical challenges. Finally, guinea pigs are an incredibly useful preclinical model for cardiac research, and as such, we have alluded to developmental similarities between guinea pigs and humans^78^. However, it is important to consider species-specific characteristics when interpreting the results of our study.

## Acknowledgements

We gratefully acknowledge Anika Haski, Anysja Roberts, Blake Cooper, and Devon Guerrelli for technical assistance related to Langendorff-perfusion studies and immunostaining experiments. We also acknowledge the Pathology and Laboratory Medicine Research Core at Children’s National Hospital for assistance with tissue embedding and slicing, and the Children’s Research Institute Light Microscopy and Image Analysis Core for microscopy access (supported by NIH grant P30HD040677).

## Abbreviations

AP: Action potential
APD: Action potential duration
CaD: Calcium transient duration
CaT: Calcium transient
CV: Conduction velocity
ECC: Excitation-contraction coupling
ECG: Electrocardiogram

